# Novel feature selection methods for construction of accurate epigenetic clocks

**DOI:** 10.1101/2022.02.21.481326

**Authors:** Adam Li, Alice E Kane, Amber Mueller, Brad English, Anthony Arena, Daniel Vera, David A Sinclair

## Abstract

Epigenetic clocks allow the accurate prediction of age based on the methylation status of specific CpG sites in a variety of tissues. These predictive models can be used to distinguish the biological age of an organism from its chronological age, and are a powerful tool to measure the effectiveness of aging interventions. There is a growing need for methods to efficiently construct epigenetic clocks. The most common approach is to create clocks using elastic net regression modelling of all measured CpG sites, without first identifying specific features or CpGs of interest. The addition of feature selection approaches provides the opportunity to reduce the cost and time of clock development by decreasing the number of CpG sites included in clocks. Here, we apply both classic feature selection methods and novel combinatorial methods to the development of epigenetic clocks. We perform feature selection on the human whole blood methylation dataset of ∼470,000 CpG features published by Hannum and colleagues (2015). We develop clocks to predict age, using a variety of feature selection approaches, and all clocks have R2 correlation scores of greater than 0.73. The most predictive clock uses 35 CpG sites for a R2 correlation score of 0.87. The five most frequent sites across all clocks are also modelled to build a clock with a R2 correlation score of 0.83. These two clocks are validated on two external datasets where they maintain excellent predictive accuracy and outperform Hannum et al’s model in accuracy of age prediction despite using significantly less CpGs. We also identify the associated gene regulatory regions of these CpG sites, which may be possible targets for future aging studies. These novel feature selection algorithms will lower the number of sites needed to be sequenced to build clocks and allow conventionally expensive aging epigenetic studies to cost a fraction of what it would normally.

## Introduction

Epigenetic clocks allow for the prediction and observation of biological aging (Bocklandt, 2011). By profiling the methylation levels at specific sites in DNA, it is possible to accurately predict the age of organisms and tissues (Horvath 2013). This is often referred to as epigenetic or DNA methylation (DNAm) age. CpG sites are areas of repetitive DNA bases where a guanine follows a cytosine, which can be modified via DNA methylation and demethylation to alter the structure of chromatin and gene expression in a cell (Moore et al., 2021). Epigenetic clocks can now predict age across multiple species and tissue types (Thompson et al., 2018), and even predict mortality (Lu, Quach et al 2019). With the increased use of DNA methylation clocks to determine biological age and screen for interventions that slow or reverse aging the demand for more robust, accurate clocks is growing.

The first epigenetic clocks were created by Bocklandt et al (Bocklandt et al, 2011) and quickly followed by the Hannum and Horvath labs in 2013 (Hannum et al, 2013; Horvath 2013). The Hannum clock, based on methylation analysis of DNA from peripheral blood mononuclear cells, was developed using elastic net regression modelling. 71 markers were selected from over 470,000 CpG sites to derive an age prediction accuracy of four years (Hanuum et al. 2013). Horvath’s clock encompasses multiple tissue types and includes 353 CpG sites that strongly predict age (Horvath 2013). Recently, there has been a focus on creating clocks with fewer CpG sites to enable epigenetic age profiling without the use of costly microarrays or expensive reduced-representation bisulfite sequencing (Ito el al. 2018, Park JL, et al. 2016, Zbieć-Piekarska et al. 2014, Spólnicka, M. et al. 2017). Alghanim et al.’s clock, built on blood methylation data, only uses CpG sites from three gene regions to explain 84-85% of age variance (Alghanim et al. 2017), and Weidner’s clock based on only 3 CpG sites, is able to predict age with an error of less than five years (Weidner et al., 2014).

Few epigenetic clock studies employ a discrete step to find optimal features for building clocks. Feature selection is commonly used in situations where the number of features far outnumber the number of samples (Guyon et al. 2003). Given the vast number of CpG sites in the genome and the relatively low number of samples in most studies, feature selection methods will improve the efficiency of clock building. Currently, the most common approach for clock building is to use a ‘correlation-with-age’ method, where CpGs that have a non-zero coefficient in ElasticNet Regression analyses are given more predictive power in the model (Horvath 2013, Hannum et al. 2013). Some clocks utilize more advanced feature selection methods such as Boruta (Renner et al., 2013), recursive feature selection (Wang et al., 2018; Darst, Malecki and Engelman, 2018; Meng, Murrelle and Li, 2008) or neural networks (Spólnicka et al. 2017). These algorithms select even fewer CpG features whilst still accurately predicting age. However the number of clocks being built with these tools is minimal and there is more room to optimise feature selection methods and parameters.

Here, we use several feature selection approaches to construct accurate epigenetic clocks with low numbers of CpG sites on the publicly available Hannum dataset (GSE40279), and evaluate their accuracy and generalizability on other datasets: GSE52588 (Horvath et al, 2015), GSE137688 (McEwen 2019), GSE85311 (Martens et al, 2020). We use a combination of modified standard methods that are readily available in python packages as well as the development of our own novel selection methods. We combine methods and use them in tandem to form new methods of feature selection, and optimise the development of epigenetic clocks to predict age.

## Results

In order to test how few CpG sites could be selected while retaining predictive accuracy, we applied each of our feature selection methods to the Hannum methylation dataset (GSE40279). Table 1 and Figure 1 summarise the results of the feature selection approaches, including the number of CpG sites identified with each approach, and the correlation (r2) with chronological age on a test set. The best model for age prediction for this dataset is *SelectKBest for 2000 features followed by Boruta*. This approach selects 35 CpG sites, with an r2 of 0.873 and a median absolute error of 3.08 years (Table 1).

**Table 1.** Results from feature selection methodology (in descending order of correlation scores).

|  | Average R2<br>Score (from<br>10CV) | STD (Years) | Mean Absolute<br>Error (Years) | Median<br>Absolute Error<br>(Years) |
| --- | --- | --- | --- | --- |
| KBest 2000 de novo then Boruta (35) | 0.873 | 0.05 | 3.82 | 3.08 |
| Intersection of all methods per CV fold then<br>Boruta (102) | 0.865 | 0.06 | 3.9 | 3 |
| KBest 25 de novo (36) | 0.862 | 0.06 | 3.96 | 3.14 |
| Boruta de novo (53) | 0.861 | 0.06 | 3.95 | 3.08 |
| %-RFE de novo to 1500 then Boruta (52) | 0.835 | 0.07 | 4.35 | 3.57 |
| ElasticNet de novo/No Feature Selection (276) | 0.827 | 0.06 | 4.64 | 3.91 |
| %-RFE de novo to 100 (161) | 0.825 | 0.07 | 4.69 | 3.83 |
| Top 10 Most Frequent (10) | 0.825 | 0.08 | 4.59 | 3.7 |
| Top 5 Most Frequent (5) | 0.82 | 0.08 | 4.6 | 3.79 |
| %-RFE de novo to 10000 then Genetic<br>Algorithm (54) | 0.818 | 0.08 | 4.61 | 3.76 |
| SFM ElasticNet de novo then Boruta (7) | 0.813 | 0.07 | 4.7 | 3.71 |
| Genetic Algorithm de novo (85) | 0.812 | 0.08 | 4.72 | 3.68 |
| SFM ElasticNet de novo (16) | 0.81 | 0.07 | 4.74 | 3.84 |
| %-RFE de novo to 1500 then SFM (16) | 0.81 | 0.07 | 4.74 | 3.84 |
| SFM ExtraTrees de novo (5) | 0.77 | 0.08 | 5.36 | 4.27 |
| SFM ExtraTrees de novo then Boruta (5) | 0.77 | 0.08 | 5.36 | 4.271 |
| Neural Network feature selection (65) | 0.76 | 0.08 | 5.65 | 4.79 |
| Post Feature Selection Intersection of all methods (1) | 0.73 | 0.09 | 5.75 | 4.38 |
| Variance Threshold de novo (2) | 0.02 | 0.02 | 11.9 | 10.61 |

**Figure 1.**
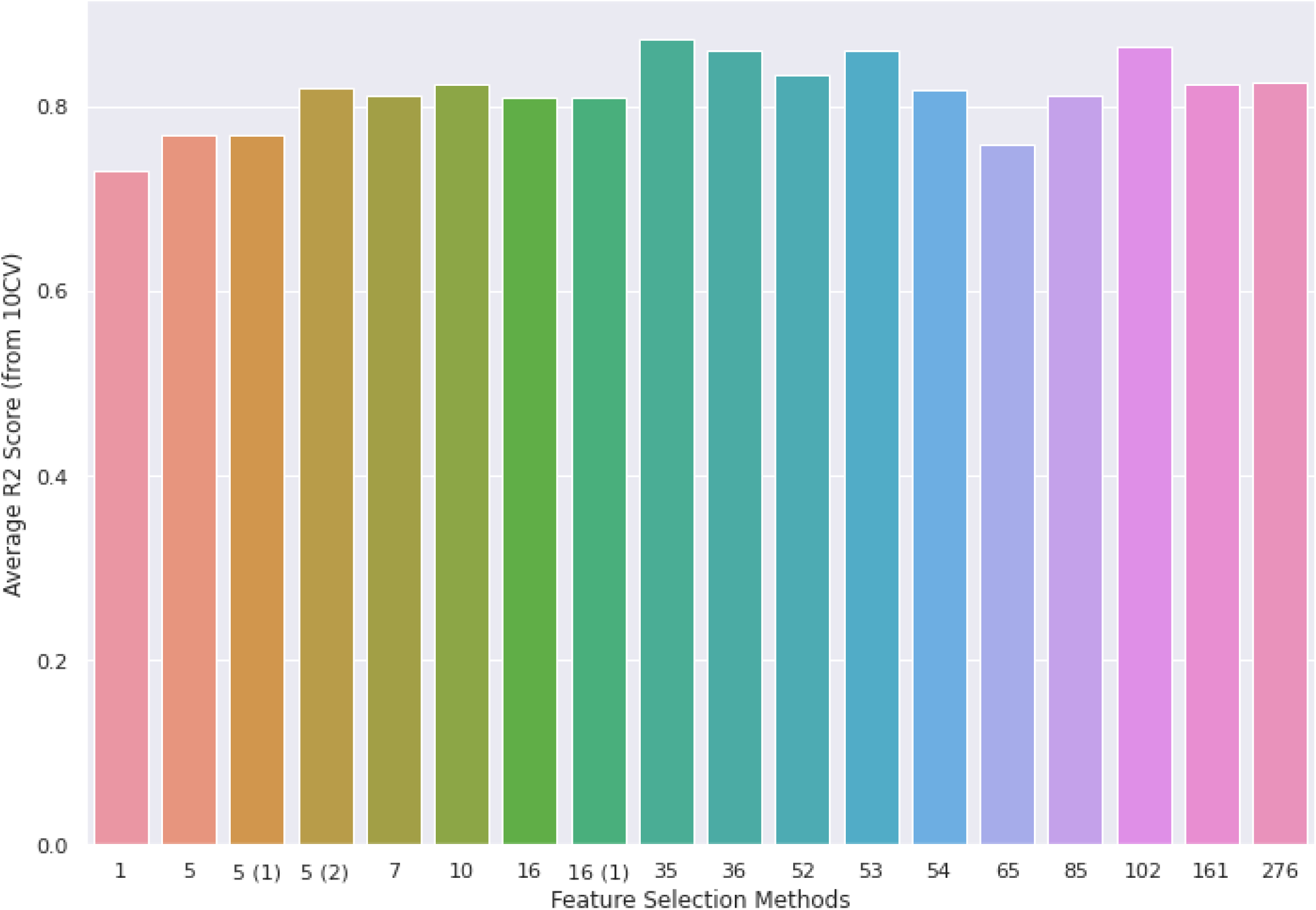
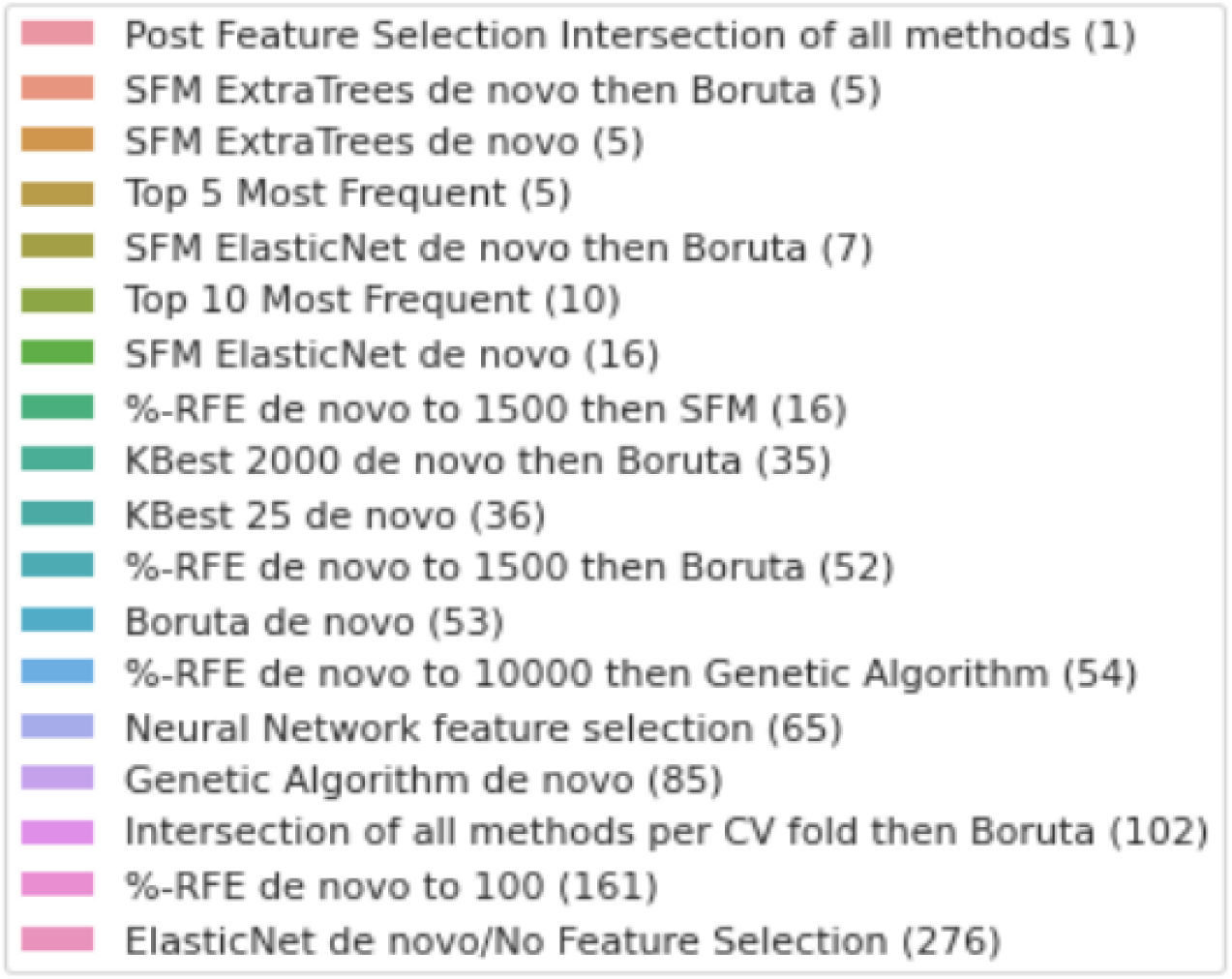
Figure showing the comparative methods with the number of features used in each model on the x-axis and their average R2 scores on the y-axis. R2 scores are relatively similar across the board despite the number of features needed for prediction varying widely.

Our other feature selection methods, including most of the SelectFromModel (SFM) methods, the genetic algorithms and several combinations of methods, achieve an accuracy of between 0.77 to 0.82 (Table 1). Despite being fundamentally different in their approach, these methods accomplish similar results and plateau in the same range of scores (Figure 1). Further optimization of each of these methods is needed to warrant their usage over other more successful methods.

*ElasticNet de novo* (Table 1, Figure 1) represents a model without any feature selection methods for comparison to the other models. This model uses all ∼450,000 features to train a model without any pre-selection or iterative algorithms. The resulting clock from this approach is based on 276 CpGs, which is a much higher number of CpGs than clocks developed with the feature selection methods (Table 1), and with a lower r2 score than five of the feature selection models (Table 1).

The majority of Recursive Feature Elimination (RFE) and Boruta based methods score 0.82 or higher suggesting that for this dataset these methods work best (Table 1). Boruta de novo and KBest 25 de novo score remarkably well with no prior method being applied (0.861 and 0.862 respectively). These are the best performing solo feature selection methods.

Using the five most frequently-selected CpGs among all the methods to build a clock gave a correlation score of 0.83 and median absolute error of 3.79 years (Table 1). Table 2 shows the corresponding GeneIDs for these CpGs. The most frequent CpG site is cg16867657 (ELOVL2) and training a clock on this single feature results in a correlation score of 0.73 (Table 1). Overall, these results demonstrate that using feature selection methods accurate epigenetic clocks can be constructed with only a few CpGs.

**Table 2.**
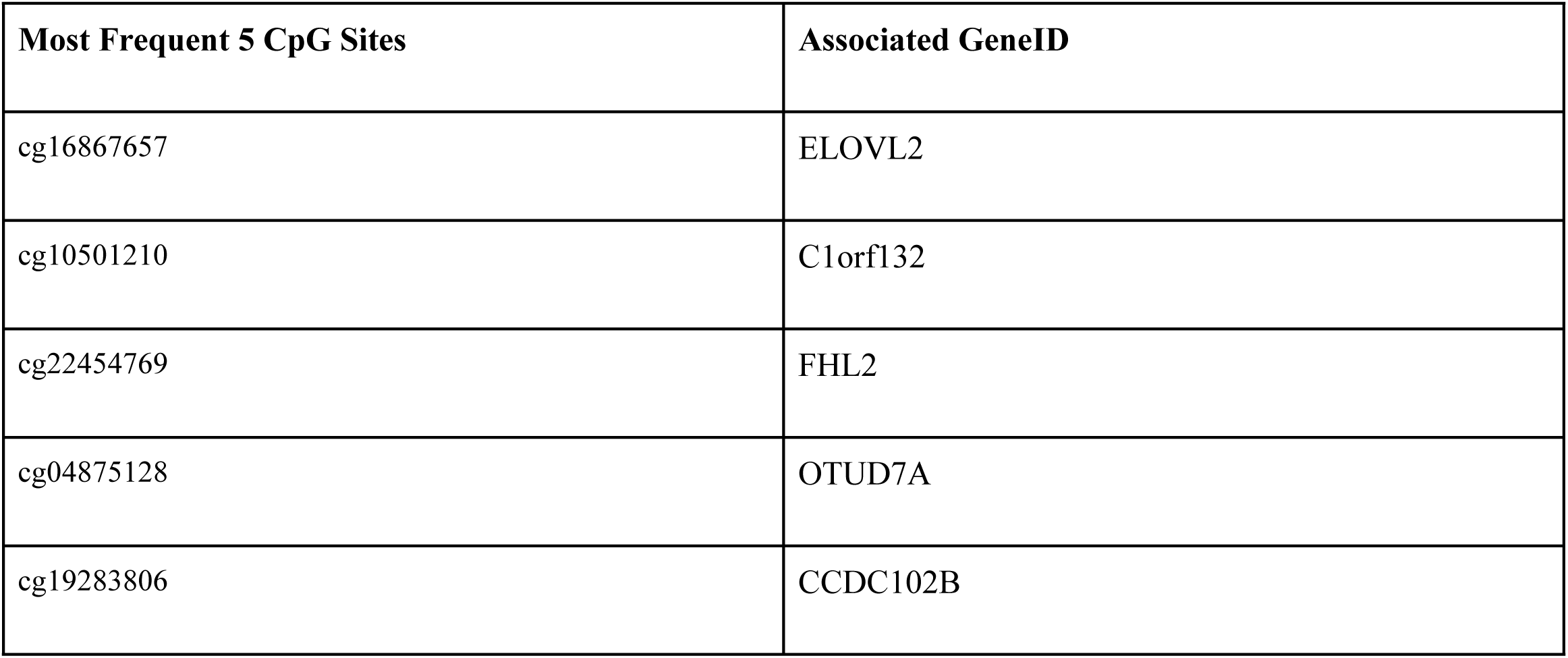
Table showing the 5 CpG sites that are chosen as most frequent predictors for aging and their associated gene symbols

| Most Frequent 5 CpG Sites | Associated GeneID |
| --- | --- |
| cg16867657 | ELOVL2 |
| cg10501210 | C1orf132 |
| cg22454769 | FHL2 |
| cg04875128 | OTUD7A |
| cg19283806 | CCDC102B |

We also tested a neural network approach for feature selection. An ElasticNet Regression model trained on the top 65 features selected by the neural network, has a moderate r2 value of 0.76. Interestingly, only four of the 65 identified neural network CpGs overlap with the CpGs selected by other methods described here. Given a neural network’s unique ability to detect these CpG sites as predictors, this is a promising predictive tool to uncover more obscure CpGs that most conventional methods miss.

We selected two models developed above for further validation of their accuracy in independent datasets. *SelectKBest for 2000 features followed by Boruta* & the *top 5 most frequent features* are the best performing feature selection method and the clock with the lowest number of CpGs sets, respectively. We applied these two clock models to two published blood methylation datasets. GSE85311 is methylation profiling of blood taken from young and old human subjects of varying exercise level (Martens et al, 2020). GSE52588 is methylation profiling of blood taken from subjects with and without down syndrome (Horvath et al, 2015). Each of the clocks predicted age very well in these external data sets with R2 values greater than 0.93 (Table 3, Figure 2), performing better than Hannum’s final published clock created from 71 CpG sites (Hannum 2013), despite using far fewer features.

**Table 3.** Table showing the results of the two final models trained on the Hannum dataset (GSE40279, Hannum et al 2013) validated on external datasets: Horvath down syndrome blood dataset (GSE52588, Horvath et al, 2015), Martens exercise blood dataset (GSE85311, Martens et al, 2020), and buccal dataset (GSE137688, McEwen 2019). Number of CpG sites/features in parentheses.

| Feature Selection Methods | Data set | r2 Score | Mean Absolute Error (Years) | Median Absolute Error (Years) |
| --- | --- | --- | --- | --- |
| KBest 2000 de novo then Boruta (35) | GSE85311 | 0.931 | 4.66 | 4.18 |
|  | GSE52588 | 0.946 | 3.35 | 2.68 |
|  | GSE137688 | 0.710 | 2.0 | 1.6 |
| Top 5 Most Frequent (5) | GSE85311 | 0.964 | 5.71 | 5.60 |
|  | GSE52588 | 0.932 | 4.56 | 3.98 |
|  | GSE137688 | 0.470 | 2.72 | 2.29 |

**Figure 2.**
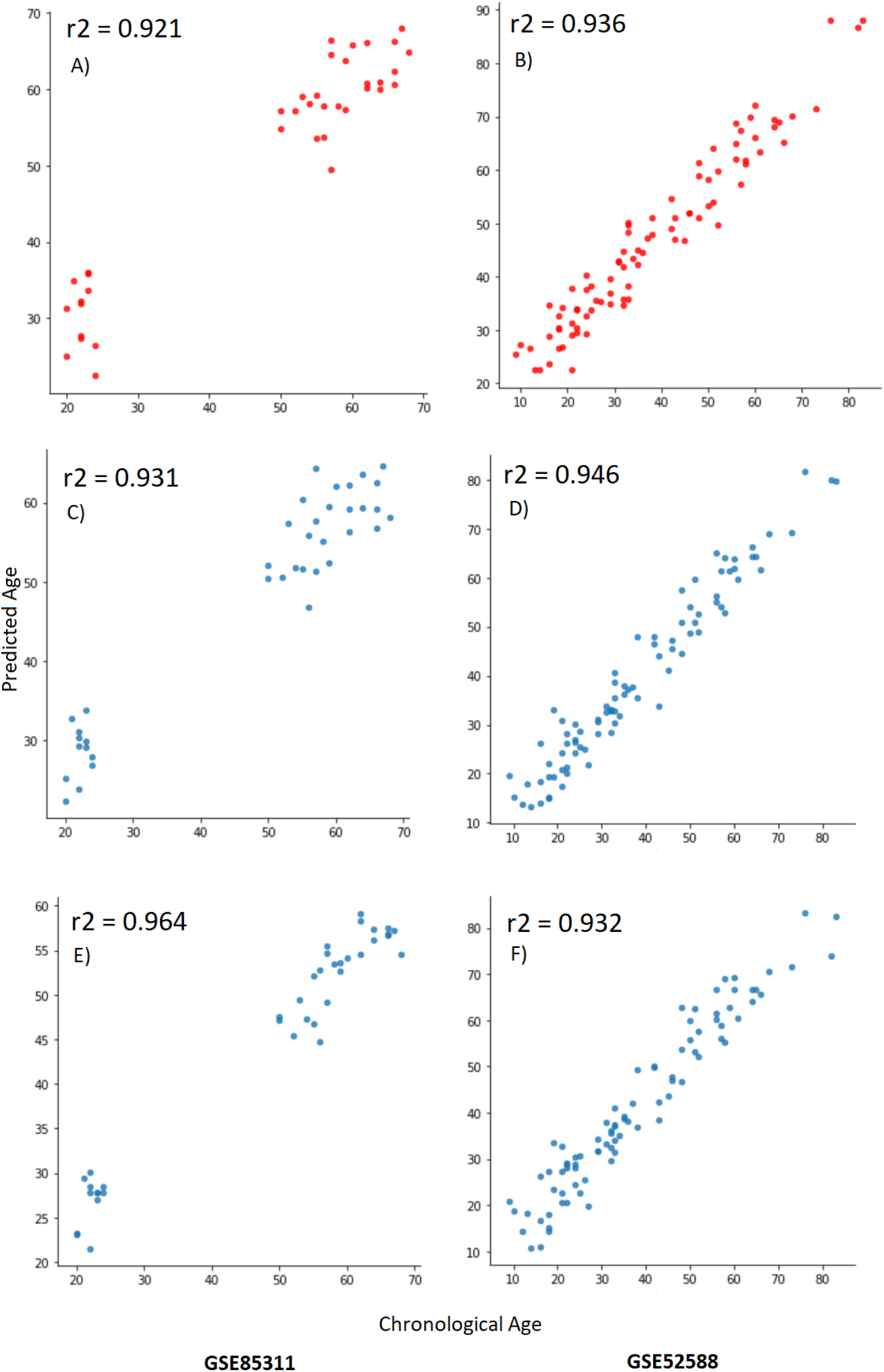
Figure showing the Predicted Ages vs Chronological Ages results of the two final models and Hannum’s model on the two external validation datasets GSE85311 and GSE52588. (A-B) Hannum (C-D) KBest 2000 de novo then Boruta (E-F) Top 5 Most Frequent.

To test whether clocks developed with our feature selection approaches can be applied to datasets other than those developed in blood, the two selected models above were also applied to a buccal cell dataset (GSE137688, (McEwen 2019)). Using the methods on this dataset, we achieved a top r2 score of 0.71 with the *SelectKBest for 2000 features followed by Boruta* method and r2 of 0.47 with the *Top 5 Most Frequent* method (Table 3). The scores were expectedly lower than the results of the other two validation sets because the clocks were trained on blood data, and applied to buccal swab data, which have inherent sampling and variance differences. While the r2 scores were not as high, the models did have very low mean and median absolute errors; the lowest of all results in this paper. Given the abundance and ease of access buccal samples provide, this is promising rudimentary groundwork for the application of feature selection methods on sample types beyond blood.

We next wanted to test whether the features selected with our methods, could be used to make accurate clocks in other datasets. We took the CpGs selected from the Hannum dataset using our top two models (SelectKBest for 2000 followed by Boruta & the top 5 most frequent CpG features), and selected those same CpGs in the Horvath down syndrome dataset (GSE52588, Horvath et al, 2015). Using only those CpGs, we created a clock from that remaining dataset, using the same cross-validation scheme (see Methods) used for the original Hannum experiment above. Remarkably, clocks developed in this dataset based on 35 features (SelectKBest for 2000 features followed by Boruta) and 5 features (top 5 most frequent) achieved r2 scores of 0.928 and 0.911 respectively (Table 4). This shows these CpGs can be used across datasets to create accurate clocks and are possibly universal, non-dataset specific CpGs for predicting age.

**Table 4.** Table showing the results of the two models created from the Horvath down syndrome blood dataset (GSE52588, Horvath et al, 2015) using the same CpGs selected from the two feature selection methods from the initial Hannum experiment. These models were validated using the same 10CV scheme from the initial Hannum experiment. Number of CpG sites/features in parentheses.

| Feature Selection<br>Method CpGs used | Average R2 Score (from<br>10CV) | Mean Absolute Error<br>(Years) | Median Absolute Error<br>(Years) |
| --- | --- | --- | --- |
| KBest 2000 de novo then<br>Boruta (35) | 0.928 | 3.39 | 2.92 |
| Top 5 Most Frequent (5) | 0.911 | 4.02 | 3.72 |

## Discussion

Overall we show feature selection methods can select CpG sites that are highly predictive of age, allowing for less features needed to build an accurate epigenetic clock. Many different types of feature selection methods are able to attain a reasonably high correlation score of around 0.75-0.85 whilst using a low number of CpG features. The rudimentary base code that outlines most of the feature selection ideas in this paper is publicly available and we hope that feature selection becomes a standard discrete step in future epigenetic clock studies. The corresponding genes of the most common CpG sites in these clocks are possible future targets for aging studies.

Two of our clocks, both trained on the original Hannum dataset, also performed well on two external datasets. The models, in fact, performed higher on validation datasets than the training dataset, and outperformed Hannum’s original clock that uses 71 features. This validates both the feature selection methods’ ability to reliably select good CpGs and the construction of our clocks. These clocks are thus able to be used by others reliably to serve as predictors of chronological age. We also applied these models to a dataset of a different sample type; buccal epithelial cells. Although the r2 scores were only moderate for this dataset, the mean and median absolute errors were the lowest we observed. This suggests an interesting future potential for buccal/saliva methylation samples, as they are much more accessible and less expensive to obtain.

In addition to the validation of the clocks, we also tested whether the identified CpGs of two of these methods could be used to make accurate clocks using the Horvath down syndrome dataset (Horvath et al, 2015, GSE52588). These clocks still achieved high 0.91-0.92 r2 scores (Table 4). This suggests that these features and their ability to predict age are not dataset specific and can universally be used across other methylation datasets.

We identified five CpGs and their corresponding genes that were of particular interest, as they were most commonly identified across all feature selection methods in our study (Table 3). Four of these CpG sites, and particularly ELOVL2, have been previously identified as strong predictors of age. ELOVL2, C1orf132, FHL2 and CCDC102B are included in an online seven CpG site epigenetic clock from the University of Santiago de Compostela (Mathgene, 2021). Zbieć-Piekarska et al constructed a linear regression model using only ELOVL2’s CpG site (cg16867657) to predict age (Zbieć-Piekarska et al., 2015) and obtained a high degree of accuracy in blood samples from humans. By manipulating the expression of ELOVL2 and observing age-related changes in the eyes of mice, Chen et al suggest that the gene is a molecular regulator of aging in the retina. Spólnicka and colleagues used ELOVL2 to accurately detect age differences from 3 disease groups (Spólnicka et al., 2018), and also highlight C1orf132 and FHL2 as key genes from which CpG sites are used for their epigenetic clock. CCDC102B also has links to aging and age-related degenerative diseases (Hosoda et al., 2018, Xia et al., 2018). Ito and colleagues developed a clock using only the CpG sites associated with CCDC102B and ELOVL2 (Ito et al., 2018) and are able to predict age with an r2 of 0.75. Additionally, Fleckhaus et al.’s study develops a clock using 8 target regions, four of which are ELOVL2, FHL2, CCDC102B and C1orf132 (Fleckhaus etc al, 2020). These papers show that our feature selection methods are able to select the most age-predictive CpG sites, consistently with other studies.

OTUD7A is the fifth gene of interest that we identified with our methods, but is the least documented. One study has previously identified that high methylation rates of CpG sites associated with OTUD7A are correlated with age (Tharakan et al., 2020), and Yin et al. identified it as a potential regulator for neurodevelopmental disorders (Yin et al.,2018). The role of OTUD7A in aging, if any, is not well-known and should be explored further.

We also applied a neural network method for feature selection in this study, but found it was not as powerful in terms of predictive accuracy as the other feature selection methods. However, this method did select many CpG features that were missed by our other conventional and novel methods. As neural network architecture becomes more advanced in its ability to read in larger datasets, the features it selects may eventually rival the accuracy of other methods. The features identified with neural networks may also give rise to new sets of CpG sites and genes worthy of study in aging.

The feature selection methods we introduce here overcome the common computational issues of stock selection methods and select a low-number of CpG sites whilst still yielding predictions of age that have high accuracy. These methods can be applied to a range of future studies developing epigenetic clocks including across new tissue types, or by examining a limited subset of CpGs in mutual overlap between bulk methylation and single cell datasets (Trapp et al, 2021). Parallelized, highly cost-reduced methods targeting specific CpG regions (Griffin et al. 2021) are another prime example. Lastly, these methods are not limited to the identification of CpG sites as features, and this pipeline could be used to identify features for biomarkers or clocks developed from a range of datasets (eg. metabolomics, microbiome, clinical data), and to predict a variety of age and health outcomes.

## Methods

### Data

The datasets for this study are from the Gene Expression Omnibus database under the accession codes GSE40279, GSE85311, GSE52588 and GSE137688. The main dataset GSE40279 we test the feature selection methods on contains 656 samples (instances) of whole blood human methylation levels at 473,035 CpG Sites (features), matched to chronological ages. All analysis was done in Python 3. All related code outlining our methods is available on github (https://github.com/adamyli/CLK-MKR).

### Cross-validation and overall approach

The main workflow methodology is outlined in Figure 3. The original dataset was split into 10 folds for cross-validation (CV). For each set of training folds, every different feature selection method was performed to select the optimal features within that training data. For every CV iteration, the intersection of each feature selection method was also recorded and we performed Boruta on the intersected features. For each of the feature selection methods, the resulting unique features from each of the 10 iterations were collected into an aggregated list and entered into a final results dataframe. This dataframe contains every unique feature selected by each selection method at each of the 10 iterations.

**Figure 3.**
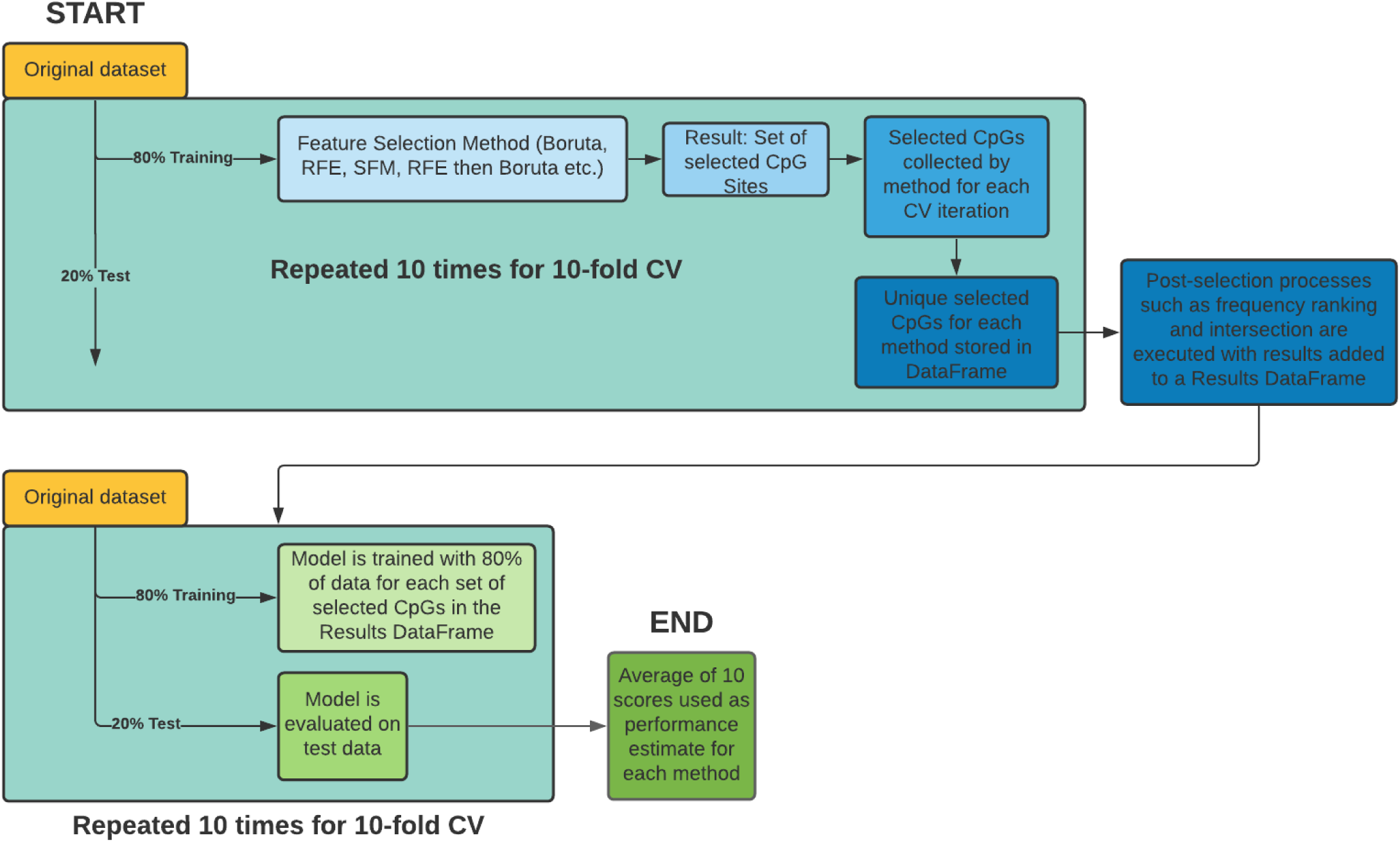
The workflow for feature selection and model evaluation. Feature selection was performed on training data for each iteration of 10-fold cross validation. The selected features of each iteration are aggregated into a list for each feature selection method type. The unique selected features for each method are collected into a dataframe where post-selection processes such as intersections are performed. We add the results to a dataframe. Each column of selected features in the results dataframe (each representing a different feature selection method) is tested using another training-testing split on the original data. This is done 10 times for 10-CV with the average of all scores being the performance estimate for that feature selection method.

Post-feature selection processes were then performed. These include the intersection between the results of all selection methods and ranking the top 5 and 10 most common features out of all the results. The results from these two post-feature selection processes were also added to the Results Dataframe. The original dataset was split into 10 folds again and for each column of the Results DataFrame, which represents the unique selected features for every method, we reduced the dataset down to the selected features. We trained the ElasticNet regression model for chronological age using training data (80%) and evaluated the model on the test data (20%) using the r2 scoring metric. For each column the mean of the 10 r2 scores was the performance estimate of that feature selection method.

The best performing model was the clock from the *SelectKBest method down to 2000 features followed by Boruta* resulting in 35 selected features. The second model of interest uses the top 5 most frequently selected CpGs. These 2 models were validated using two external blood methylation datasets; (GSE52588) and (GSE85311) and their performance was compared to Hannum’s model’s predictions on these two datasets. The features from these two models were also used to build models from the GSE52588 dataset and predict age using the same 10-fold CV as the Hannum dataset to investigate if these selected features are effective across datasets.

These two models are also applied to a methylation dataset taken from buccal cells (GSE137688) to see if performance could be replicated in conventionally cheaper samples (McEwen 2019).

### Feature selection methods

#### SelectFromModel (SFM)

SFM is a function within skLearn (Pedregos et al., 2011) that wraps around and trains a model on a dataset and allows the user to specify a threshold of feature importance. Depending on whether the model is a standard regression or random forest model, the feature importance is calculated from the coefficients or mean importance respectively. Features (CpG sites) with less feature importance than this threshold are discarded, leaving only the features with the highest coefficient or importance. This method is fast but simple. Thresholds of 0.01, 0.05, 0.1, 0.5 are tested. For this study, the models that the SFM wraps around are ElasticNet Regression and ExtraTrees forest.

The ExtraTrees Regression estimator is composed of a number of decision trees. A decision tree can be thought of as an intuitive flowchart where an answer to one decision between 2 or more choices leads to another. Decision trees decide how to split by prioritizing the split that creates the least uniform distribution of labels or values. This branching of nodes continues until it reaches a node that cannot decide which split to use because they result in equally uniform distribu-tions - meaning any more branches will not help the tree make any better decisions. In this sense ExtraTrees is similar to the more popular random forest with a few distinct differences. Random Forest samples the training data with replacement to train their decision trees whileExtraTrees uses the entire original dataset. However ExtraTrees randomly chooses the split instead of optimally finding a locally one which is what Random Forest does. ExtraTrees are therefore less exhaustive in their optimization and are faster than Random Forests. This is ideal for us as a Random Forest with 5-8 trees in it can take several hours to train on a dataset as large as ours. A random forest takes an advantage known as bagging by taking random instances of the dataset and training its model from solely those samples. For a regression problem like ours the average value of all trees are taken as the final prediction.

#### Recursive Feature Elimination (RFE) and the introduction of %-RFE

RFE is a function that trains a model on a dataset and removes the weakest feature based on the lowest feature importance from the dataset (Pedregos et al., 2011). This new dataset of N-1 features is trained again with a model and the process is repeated until only the user specified number of features is left. By removing 1 feature each time, RFE is a brute force algorithm that leaves only the best performing features at each iteration. However it does not take into account all features at the same time, and is unable to be aware of relationships between CpGs when it comes to predicting age e.g. some CpGs may become a strong predictor of ageing in the presence or absence of another.

Applying the stock RFE algorithm to our dataset of 473,035 features is computationally limiting due to the size of the dataset (Supplementary Table 1). Instead, we write an algorithm that removes a percentage-based number of features at each iteration allowing us to aggressively remove the majority of unnecessary features at the start but be more meticulous with our selection near the end. The percentage chosen is 1%, i.e. removing 4730 features at 473,035 and 1 feature at 100.

#### Boruta

RFE is a ‘minimal optimal’ feature selection method, meaning it attempts to select the smallest set of features with the minimum error for an estimator and aims to optimize this ratio. Boruta differs as an ‘all-relevant’ feature selection method compatible with only tree-based regression methods, such as random forests (Kursa et al., 2010). Instead of trying to find the most compact set of features to predict with, it considers all features that could possibly contribute towards prediction overcoming the weakness of RFE’s greedy nature. Boruta creates duplicates of the existing features with randomized values called ‘shadow features’. The dataset comprising the original and the shadows, is trained on the tree estimator and the shadow features compete with their original forms. Features that consistently beat their shadow counterparts are selected as reputable predictors. In order to deal with the computational power needed to train a random forest with over 470,000 features, we use fewer trees and adjusted iteration counts in these models.

#### SelectKBest

SelectKBest is a feature selection method in sklearn similar to SFM that fits a dataset and selects features based on a scoring metric (Pedregos et al., 2011). For each feature it calculates the correlation value between the feature and target label and ranks them. This method is fast due to its shallow nature of only training once so is not useful when used alone. However, it is helpful to reduce the total number of features for usage of more greedy algorithms such as Boruta. In our methodology we select the top 25 features and the top 2000 features using SelectKBest. We perform Boruta on the top 2000 features.

#### Variance Threshold

Variance threshold is a simple and exploratory method that removes all features whose column of values do not reach the threshold of variance (Pedregos et al., 2011). Since some datasets naturally may not have a high degree of variance in their recorded data, this method is not consistent. However since its execution is the fastest out of all the methods (Supplementary Table 1) it is included as an added method.

#### Neural Network (NN) Feature Selection

The rudimentary neural network is built using PyTorch to feature select CpG sites, as neural networks have been known to capture nonlinear relationships between data points. We were interested in seeing what would be good predictors of aging that might have been missed by the other linear regression models and lay the groundwork for future feature selection using NNs. As a proof-of-concept we used %-RFE to reduce the number of features from 473,035 down to 100. The NN first uses all 100 original features and trains the model once, its score being recorded as a benchmark. Following this, for each of the 100 features, the NN is then trained twice; once where all methylation levels of that feature equals 1 and once where they all equal 0 to simulate the CpG being fully methylated and also absent. Both are done to account for the cases where the original methylation value is close to 0 or 1. The mean of the two resulting scores are compared to the benchmark with the difference being recorded for each CpG site. The CpG sites are ranked in difference to establish an idea of feature importance with the postulation that a larger difference between the presence and absence of the CpG will insinuate that the CpG has a greater impact on age prediction. The top 50-75 are recorded as selected features.

#### Genetic Algorithm

An algorithm based on the nature of Darwinism evolution where a population of ‘creatures’ are assigned a desired amount of features from the original dataset at random. These creatures are evaluated via predicting a validation set and assigned a score or ‘fitness’. The lowest scoring creatures are culled next, simulating survival of the fittest. The remaining creatures are bred by creating a child creature that has features from their shared ‘gene pool’ and having a new number of them selected randomly. There is a chance for a certain number of these ‘genes’ to be mutated. Meaning some of the features will be randomly swapped for a different one from the original dataset. This helps introduce variation. This process is repeated for a specified number of generations or until a desired fitness is met.

The genetic algorithm is powerful as it allows the user many points of optimization, depending on the creativity of the user. For instance, the number of generations, number of features and creatures are all linked variables where a perfect balance can be found. When it comes to the breeding process it is possible to implement a ‘polygamous’ aspect where a highly successful creature is allowed to breed multiple times to ensure the most predictive features are passed on and tested further in other combinations. Mutation rate, number of genes allowed to mutate as well as number of children produced per breed (with possibility of scaling number of children produced with the fitness of the parent). It is also common for genetic algorithms to be run in parallel, predicting subsets of a label, e.g. an algorithm for young samples and one for old.

#### Novel methods combining multiple feature selection methods

The introduction of %-RFE and to a lesser extent SFM allows us to synthesize novel feature selection methods. %-RFE allows for the removal of ‘fluff’ down to a more manageable number of features (usually a few thousand) and allows for more powerful methods to be used such as Boruta, Neural Networks and RFECV. These methods require more iterations and computational power so being able to distill down to the most important thousand features to choose from is ideal. The synthesized methods consist of %-RFE first selecting features to an amount appropriate for the next method. SFM is also used as a preliminary selection method in this way. The final synthesized methods consist of modular code functions that allow us to alternate the order in which the selection methods are used as well as let us combine them together and use the output of onemethod as the input of another.

#### Clock Models

The epigenetic clocks are built using ElasticNetRegression models. ElasticNet is chosen as it is the current standard for epigenetic clocks and outperforms Random Forests and SVMs with these data and feature selection methods.

This model is a variant of classical linear regression. This aims to solve for the coefficients of a linear equation that equals the ‘best fit line’. The best fit line minimizes the sum of squares by having the least distance between the data points and the line. The equation for ordinary linear regression is as follows:

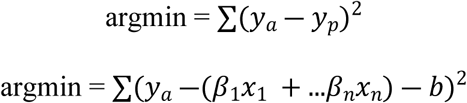

Where y_a is the actual value of the target label and prediction y_p calculated by the summation of predictors ‘x’ multiplied by a vector of coefficients β_n that is found from fitting the model b.is the y-intercept. argmin signifies a cost function where we seek to minimize the answer given input arguments.

Regularization is a process in which different variants of bias and penalties are introduced to assist in finding the solution to this equation that allows for the best predictive accuracy. These penalties are controlled by a lambda value (alpha in sklearn) that controls how heavy (large) this penalty is. The L1 penalty is referred to as Lasso Regression, it adds a bias that is the absolute value of the coefficients. The L2 penalty is referred to as Ridge regression, this adds a bias that is the squared value of the coefficients. Unlike ridge regression, lasso regression can shrink the coefficients of unneeded parameters (features) to 0 (due to the penalty term not being squared), essentially eliminating them, leaving only useful features. Lasso can be quite aggressive however, taking only 1 feature out of several correlated ones or selecting too few. This is where ElasticNet comes in. The generic form of the ElasticNet equation is:

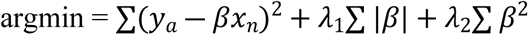

Where L1 is the regularization penalty for the ’Lasso’ part of the regression equation andL2 is the penalty for the ‘Ridge’ portion (Zou, Hastie. 2005). ElasticNet combines both Lassoand Ridge regressions, adding both terms to the equations. Each penalty gets an indepen-dent alpha / lambda that is tuned via cross-validation or other methods. This method allows the best of both worlds depending on the feature.

## Acknowledgements

D.A.S. was supported by the Glenn Foundation for Medical Research; A.E.K was supported by a K99 award from the NIH (K99 AG070102); A.M. was supported by an F32 Ruth L. Kirschstein National Research Service Award (NRSA) Individual Postdoctoral Fellowship from the NIH (F32 AG069363);

D.V. received financial support from the NIDDK Mouse Metabolic Phenotyping Centers (RRID:SCR_008997, MMPC, www.mmpc.org) under the MICROMouse Funding Program, grants DK076169, and a NIH T32 grant T32AG023480.

